# Accurate Detection of Convergent Mutations in Large Protein Alignments with ConDor

**DOI:** 10.1101/2021.06.30.450558

**Authors:** Marie Morel, Frédéric Lemoine, Anna Zhukova, Olivier Gascuel

## Abstract

Evolutionary convergences are observed at all levels, from phenotype to DNA and protein sequences, and changes at these different levels tend to be highly correlated. Notably, convergent and parallel mutations can lead to convergent changes in phenotype, such as changes in metabolism, drug resistance, and other adaptations to changing environments.

We propose a two-step approach to detect mutations under convergent evolution in protein alignments. We first select mutations that emerge more often than expected under neutral evolution and then test whether their emergences correlate with the convergent phenotype under study. The first step can be used alone when no phenotype is available, as is often the case with microorganisms. In the first step, a phylogeny is inferred from the data and used to simulate the evolution of each alignment position. These simulations are used to estimate the expected number of mutations under neutral conditions, which is compared to what is observed in the data. Next, using a comparative phylogenetic approach, we measure whether the presence of mutations occurring more often than expected correlates with the convergent phenotype.

Our method is implemented in a standalone workflow and a webserver, called ConDor. We apply ConDor to three datasets: sedges PEPC proteins, HIV reverse transcriptase and fish rhodopsin. The results show that the two components of ConDor complement each other, with an overall accuracy that compares favorably to other available tools, especially on large datasets.

## Introduction

Convergent evolution is often defined as the independent acquisition of similar traits in distinct lineages over the course of evolution (Arendt and Reznick 2008; Losos 2011; Stern 2013). The studied traits can be behavioral, morphological, molecular, etc. In each category, traits can be quantitative (size, length, weight…), binary (presence or absence of a given phenotype) or categorical (a trait is subdivided into several categories). The presence of convergence, especially at the phenotypic level, is often seen as evidence of adaptation in the sense that similar evolutionary paths were found in response to the same evolutionary constraints (Castoe et al. 2009; Losos 2011). Many studies focus on the molecular level, assuming that convergent phenotypes may result from the same genetic changes (Stern 2013; Rosenblum et al. 2014; Storz 2016). At the protein level, it is common to distinguish (Zhang and Kumar 1997) between parallel mutations (a change toward the same amino acid is observed from the same ancestral amino acid), convergent mutations (change toward the same amino acid, from different ancestral amino acids), and reversions (mutations that restore an amino acid previously lost during evolution). For the sake of simplicity, in what follows we will refer to these three types of mutations as “convergent mutations”, unless explicitly stated.

Examples of evolutionary convergence at the molecular level have been demonstrated in higher eukaryotes, related to adaptation to certain environments (Muschick et al. 2012; Foll et al. 2014; Foote et al. 2015; Hill et al. 2019; Lu et al. 2020; Xu et al. 2020), diet (Zhang 2006; Zhen et al. 2012; Ujvari et al. 2015; Hu et al. 2017), changes in metabolism (Besnard et al. 2009; Parto and Lartillot 2018), morphological transformations (Larter et al. 2018) and acquisition of new abilities (Davies et al. 2012; Parker et al. 2013; Lee et al. 2018; Marcovitz et al. 2019; Chai et al. 2020). Similarly, when submitted to constraints such as harsh experimental conditions and drug treatments, microorganisms and viruses adapt and are likely to exhibit similar escapes. This has been demonstrated in HIV after exposure to antiviral drug treatments in several patients (Crandall et al. 1999) and within a single treated patient (Holmes et al. 1992). Similarly, Cuevas et al. (2002) found adaptive convergence in experimental populations of RNA viruses, and van Ditmarsch et al. (2013) in pathogenic bacteria. In natural conditions, evolutionary convergence was found in viruses having experienced host shifts (Longdon et al. 2018; Escalera-Zamudio et al. 2020; Martin et al. 2021) and changes in vector specificity (Tsetsarkin et al. 2007).

Several methods have been developed to detect convergent evolution at the molecular level (Zhang and Kumar 1997; Zhang 2006; Tamuri et al. 2009; Parker et al. 2013; Thomas and Hahn 2015; Zou and Zhang 2015a; Parto and Lartillot 2017; Chabrol et al. 2018; Rey et al. 2018). They are all based on prior knowledge of a convergent phenotype and aim to identify the protein mutations underlying the phenotypic trait studied. However, they differ in the scale at which molecular convergence is sought and the exact definition of what a convergent mutation is.

Some approaches aim to identify which coding genes harbor mutations supporting a convergent phenotype, while others study which amino-acid changes can explain convergent changes at the scale of a single protein. Methods of the first category are commonly applied to eukaryotic and prokaryotic genomes and perform genome-wide analyses to detect convergent genes by considering simultaneously all positions of the corresponding protein sequences; for example, the methods developed by Parker et al. (2013), Zou and Zhang (2015b), Thomas and Hahn (2015) and Chabrol et al. (2018) were applied to the search of genes responsible for echolocation in mammals. In the second configuration, the coding genes responsible for the convergent phenotype have already been identified and the methods focus on the detection of convergent evolution at the position level; for example, Zhang and Kumar (1997) identified convergent and parallel mutations in stomach lysozyme sequences of foregut fermenters. Similarly, Zhang (2006) found parallel substitutions in colobine pancreatic ribonucleases, and Rey et al. (2018) found positions with convergent substitutions in the PEPC protein occurring jointly with the transition toward C4 metabolism in sedges. In fact, testing the significance of convergent changes at individual protein positions has many potential applications. In the case of complex eukaryotic and bacterial organisms, there are few examples of a single amino-acid change that could explain a convergent phenotype (Storz 2016). However, in the case of viruses with rapid evolution, and whose (small) genomes are strongly constrained, only a few amino-acid changes are generally possible at a given position (Pond et al. 2012) and position-wise convergent evolution is expected to be relatively frequent (Gutierrez et al. 2019). Determining molecular changes that deviate from what is expected by chance can thus be indicative of adaptive phenomena. This was the case for SARS-CoV-2, where one first identified mutations in the Spike protein, which were spreading within the viral population and appeared multiple times independently, before being demonstrated to be evolutionarily advantageous for the virus (van Dorp et al. 2020; Korber et al. 2020; Martin et al. 2021). Note, however, that mutations that were initially thought to be adaptive were eventually shown to be simply the result of founder events (Hodcroft et al. 2021), demonstrating the difficulty of detecting convergent mutations without access to the phenotype.

Most importantly, different methods have different ways of selecting which mutations underly the studied convergent phenotype. In the most intuitive definition, one aims to detect mutations toward the same amino acid, which occurred in all clades with the convergent phenotype. This is the definition used first in (Zhang and Kumar 1997) and then in (Zhang 2006; Foote et al. 2015; Thomas and Hahn 2015; Zou and Zhang 2015b). An extension was proposed by Chabrol et al. (2018), where the convergent amino acid may only be found in a subset of the convergent species, as well as in some non-convergent species. Considering that a change toward the same amino acid may be too strict since several amino acids have similar physicochemical properties, Rey et al. (2018) relaxed this constraint in the PCOC program, by considering changes in amino-acid profiles (Le et al. 2008). Their work on amino-acid profiles follows previous works aimed at detecting positions under condition-dependent selection, but which did not focus solely on convergent evolution (Tamuri et al. 2009; Parto and Lartillot 2017; Parto and Lartillot 2018). A radically different approach, proposed by Parker et al. (2013) and inspired from Castoe et al. (2009), relies on the fact that convergence can lead to errors in phylogenetic reconstruction by artificially bringing convergent species together. These authors proposed selecting positions that best support the phylogeny that groups species with the convergent phenotype together, rather than the species tree (but see the critiques of this method by Thomas and Hahn (2015) and Zou and Zhang (2015b)).

One of the main challenges in detecting molecular convergence is to identify only the convergent mutations that are linked to the studied convergent phenotype. In their review of methods for detecting molecular convergence, Rey et al. (2019) referred to this type of mutation as foreground convergence (or foreground convergent mutations) in opposition to background convergence which is unrelated to the convergent phenotype. Indeed, at the molecular level, one can find patterns of convergent mutations linked to another convergent phenotype, or occurring because of mutational biases, protein conformation limitations, constraints at the molecular level or epistatic forces (Zhang and Kumar 1997; Rokas and Carroll 2008; Storz 2016; Stoltzfus and McCandlish 2017). It has been shown that most (if not all) substitution models may fail at distinguishing between foreground convergent mutations and background ones (also called non-adaptative convergent mutations), especially in close taxa between highly exchangeable amino acids, and on fast-evolving sites (Goldstein et al. 2015; Zou and Zhang 2015a). In other words, finding multiple independent mutations resulting in the same (or a similar) amino acid should be tested carefully, even when the number of such mutations appears to be high. We shall see that our findings confirm this.

Another difficulty is the definition of the convergent phenotype and the annotation of taxa that do or do not have this phenotype. For example, in the case of viruses, we usually do not know the exact phenotype, but use a proxy instead. In the case of drug resistance mutations (DRMs) that occur repeatedly in different patients treated with antiviral drugs, we use the treatment status as a proxy for the resistance status. Although we expect that most (but not all, e.g., due to poor adherence) sequences from patients who fail drug treatment will contain resistance mutations, we also expect that some drug resistance mutations will be found in untreated (naive) patients in the case of resistance transmission (Blassel et al. 2021). Similarly, environmental constraints are not strictly speaking phenotypes, but act as selective forces that can lead to phenotypic and molecular convergence. However, we do not expect all organisms living under the same environmental conditions to exhibit the same recurrent mutations.

In some respects, the identification of convergent mutations has similarities with the detection of positions under positive selection (Goldman and Yang 1994). The idea is indeed to identify mutations that might be advantageous, as they are found more often than expected in a neutral (or purifying) model of evolution. In the positive selection framework, these mutations can be directed to a specific amino acid (directional), or correspond to any change that differs from the original amino acid (diversifying). This is the case, for example, with immune avoidance where mutations towards any new amino acid at antigenic sites are generally favorable and positively selected. Conversely, in the case of convergent evolution, we are interested in substitutions towards one or a few similar amino acids, in the branches leading to the convergent taxa. Thus, a large number of non-synonymous substitutions on convergent positions are expected, but the criterion of positive (or relaxed purifying) selection alone is not sufficient to assert convergence. The FADE software (Murrell et al. 2012) in the HyPhy suite (Pond et al. 2005) tests whether positions in a protein alignment are subject to directional selection (or mutational bias) within a specified set of “foreground” branches that typically correspond to convergent taxa. This tool thus has its roots in positive selection approaches, but is closely related to convergence detection.

Here we propose a new method for detecting convergent evolution at the position (or site) scale in large amino-acid alignments, while relaxing the constraint that convergent mutations must be found only in organisms with the convergent phenotype and in all of them. Our method does not require specifying the branches where molecular convergence occurred (as with PCOC and FADE, for example), which is a complex step, especially with large data sets and when using a proxy for the phenotypic convergence. The taxa are simply annotated as convergent or non-convergent, and the mutations correlated with this status are then detected. We are interested in mutations leading to a target amino acid, regardless of the ancestral amino acids at this position. In other words, parallel, convergent mutations and reversions are considered indifferently, and we consider mutations resulting in different target amino acids as different events. With this definition our method is in line with methods aiming at detecting changes towards the same amino acid, as opposed to detecting changes in profiles (Rey et al. 2018). Indeed, there are many examples of known convergent mutations, where the changes involve highly exchangeable amino acids that have very similar biochemical profiles. For example in HIV drug resistance, there are convergent mutations from Isoleucine to Valine and Tyrosine to Phenylalanine (the two most exchangeable amino acid pairs, cf. BLOSUM62) that confer resistance to certain drugs (Wensing et al. 2019).

In the following sections, we describe this approach, which is implemented in a workflow called ConDor (for Convergence Detector), available as a web service (condor.pasteur.cloud) and as a standalone workflow. Its performance is evaluated using three datasets involving sedge PEPC proteins, HIV reverse transcriptase, and fish rhodopsin. The results are compared to those of PCOC and FADE, which are based on different assumptions.

## New Approaches

### Method overview

Our method aims to identify amino-acid mutations that emerged multiple times in independent lineages, occurred more frequently than expected under a neutral (or null) substitution model, and are correlated with a known convergent phenotype. The method is subdivided into two independent components: (1) the “Emergence” component that detects mutations emerging more often than expected under neutral evolution, and (2) the “Correlation” component that identifies mutations that are positively correlated with the convergent phenotype. The combination of the two components accurately identifies amino-acid mutations resulting from convergent evolution associated with the convergent phenotype (foreground mutations), although it is also possible to execute and interpret the results of the two components independently.

A representation of the ConDor workflow is shown in Figure 1. Inputs are constituted of: (i) a multiple protein sequence alignment (MSA); (ii) a phylogeny; (iii) an outgroup; (iv) the phenotype of each of the taxa; and (v) user supplied thresholds to select convergent mutations. The quality of the MSA and phylogeny is critical and should be carefully checked by users of the method, as, for example, running ConDor on poorly aligned and gappy sites of a MSA can lead to poor results and incorrect conclusions. The two first steps of the workflow (Fig. 1) are common to the Emergence and Correlation components. In step (1), we estimate the parameters of the null model from the MSA (parameters of the substitution model, ML-based branch lengths of the phylogeny, evolutionary rate per position, etc.). In step (2), we reconstruct the substitution history and count the number of emergence events of mutation (EEMs) observed for every position and amino acid of interest in the MSA (i.e., those present in sufficient number of sequences). In the Emergence component, for each position and amino acid of interest, the two main steps are: (3) simulation of new datasets under the null model and counting of simulated EEMs; (4) comparison of observed and simulated numbers of EEM to identify the mutations that occurred significantly more often than expected assuming the null model. The Correlation component is applied to all mutations with more than *n* EEMs (*n* is user-defined). The two mains steps are: (3’) computation of the log Bayes factor of the model assuming a dependence between the presence/absence of the phenotype and the given mutation, versus the model assuming their independence, using BayesTraits (Pagel 1994; Pagel and Meade 2006); (4’) determination, for mutations associated with the phenotype, whether the dependence is positive or negative. The final step (5) combines the results of steps (4) and (4’) and provides a list of potential convergent mutations. The results of steps (4) and (4’) are also provided to the user and can be interpreted independently.

**Figure 1:**
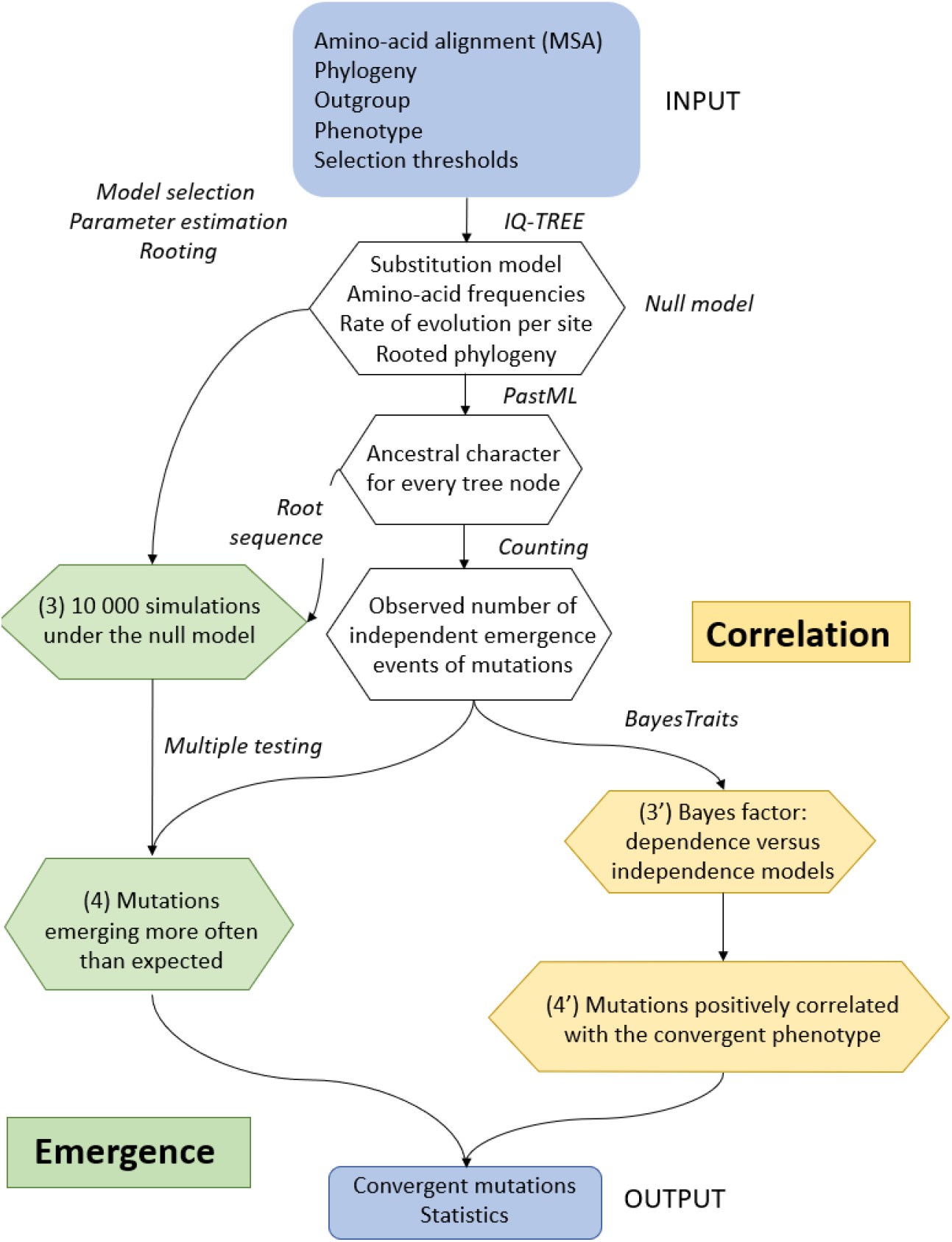
Flowchart of the method. The method takes as input an amino-acid alignment as well as the corresponding phylogeny and phenotype metadata. The MSA and phylogeny are used for inference of the null model (branch lengths, substitution model and its parameters, evolutionary rate per site, etc.) and ancestral character reconstruction. In the emergence component, the tree and root sequence are used to simulate 10,000 alignments under the null model; the output is the list of mutations that emerged more often in the input alignment than in the simulations. In the correlation component, we select mutations that are positively correlated with the phenotype. The combination of the two components gives the list of mutations proposed as convergent.

Note that step (4) does not require knowledge of the phenotype (or a proxy for it, as with DRMs in HIV). For all selected mutations, Condor provides the evolutionary rate of the corresponding position, the nature of the mutation (convergent, parallel, revertant), the number of EEMs, the genetic barrier (minimum number of mutations at the DNA level), the BLOSUM score, etc. All these statistics are described in the user guide (https://condor.pasteur.cloud/help) and can be used to further analyze the results and select the most relevant mutations.

The null model and all its components are inferred from the input alignment and the phylogeny using ModelFinder (Kalyaanamoorthy et al. 2017) and IQ-TREE (Nguyen et al. 2015). The selected substitution model, along with amino-acid frequencies, rates-across-sites distribution parameters, branch lengths, and evolutionary rate per site are assumed to represent the data without convergence. We make this assumption because using large alignments (>1000 sequences), we consider that mutations resulting from convergent evolution are rare enough to have a negligible influence on parameter inference. The phylogeny with optimized branch lengths is then rooted using the user supplied outgroup. This is necessary to infer the ancestral sequence at the root of the tree, run simulations starting from this sequence, and count simulated EEMs. For the analyses discussed here, we restricted ourselves to mutations present in at least 0.5% of the sequences (default value, this threshold is adjustable by the user). Ancestral character reconstruction (ACR) for positions with mutations of interest is performed using a maximum likelihood approach, implemented in PastML (Ishikawa et al. 2019). We use the “maximum a posteriori” (MAP) method in which the state with the highest marginal posterior is selected at each tree node. Once all ancestral amino acids are reconstructed and associated with all internal nodes in the phylogeny, we identify where independent amino-acid changes occurred in the tree and count them as explained in the subsection “Counting emergence events”. Using this count of the observed number of EEMs, we restrict the Emergence and Correlation analyses to mutations with more than *n* EEMs (i.e., found in more than *n* independent clades, *n*=2 in all analyses below).

### Estimating the expected number of emergences with simulations

The emergence component consists of simulating the convergence-free evolution that is expected for each tested position of the alignment. In our experiments, we performed many (10,000) simulations per position, using a homemade script (available on GitHub). Our implementation does not use the exact root amino acid reconstructed by ACR as a starting point, but instead draws amino acids based on their marginal posterior probabilities to account for reconstruction uncertainty (e.g., two amino acids with posteriors of 0.55 and 0.45). After the simulations of sequence evolution along the tree, we count the number of EEMs (10 000 values per position and per AA studied) using the algorithm detailed below. For example, let us consider the mutation M41L from our HIV dataset, where at position 41, a Methionine (M) is substituted by a Leucine (L) in 211 sequences. The observed number of EEMs toward L is 47, which is smaller than 211 as in some subtrees all tips have L, corresponding to only 1 EEM. Then, 47 is compared to the distribution of the number of EEMs toward L (always starting from an M at the tree root since there is no ambiguity in ACR), among 10,000 simulations in the null model; this distribution ranges from 0 to 31 with an average of 12. From the observed number of EEMs and the distribution of simulated EEMs, we estimate a p-value. To avoid zero p-values when all simulations result in fewer EEMs than observed EEMs, we use a pseudo-count of 0.5, which means that the (uncorrected) p-value is equal to (0.5 + #simulated-EEMs ≥ observed-EEMs) / 10,001 (∼5×10^−5^ in our M41L example). Since we test many positions and mutations, we use the Holm–Bonferroni method (Holm 1979) to correct for multiple comparisons, with a default rejection threshold of 10% (adjustable by the user). We consider that mutations passing threshold after p-value correction did not occur by chance. These mutations can be studied on their own in the absence of an identified phenotype. However, we know from previous studies that background convergent mutations in real data tend to be more frequent than expected under any available substitution model, due to model approximations, epistatic constraints, etc. (Rokas and Carroll 2008; Castoe et al. 2009; Goldstein et al. 2015; Zou and Zhang 2015a). Moreover, some of these very frequent mutations may be truly adaptive and convergent, but for other phenotypic traits than the one studied. Thus, we expect that a significant fraction of these frequent mutations are false positives for the studied phenotype. The Correlation component complements the Emergence component to focus on mutations that correlate positively with the phenotype, that is, foreground convergent mutations.

### Counting independent emergence events of mutation (EEMs)

To count EEMs, we consider only the initial appearances of each mutation in the tree. However, the observed number of EEMs is inferred by ACR from the input sequences, while the expected number of EEMs and its distribution are estimated from many simulations evolving the probabilistic root sequence along the tree using the null model. In simulations, there may be changes in internal nodes that are not transmitted to any tree leaf. In this case, these changes cannot be inferred by ACR, and the expected number of changes artificially deviates from the ACR-based observed number of EEMs. This effect is even more pronounced on fast-evolving positions since more changes are expected. For this reason, we decided to only count as EEMs the changes transmitted to at least one leaf.

In the tree illustrated in Figure 2, there are 6 changes along the branches, which are represented either by a cross or a NO symbol. The NO symbol stands for a change that is not transmitted to any tree leaf (blue to red on the bottom part of the tree). This node would have been inferred either blue or yellow by ACR, but highly unlikely red, while this might occur in simulations. This is why we do not count these cases. The two yellow crosses mark changes that are transmitted to at least one leaf, and are both counted. The two blue crosses mark a reversion to the root state. Even though we do not make the difference between convergent, parallel, and reversion events, we keep the information for downstream interpretation.

**Figure 2:**
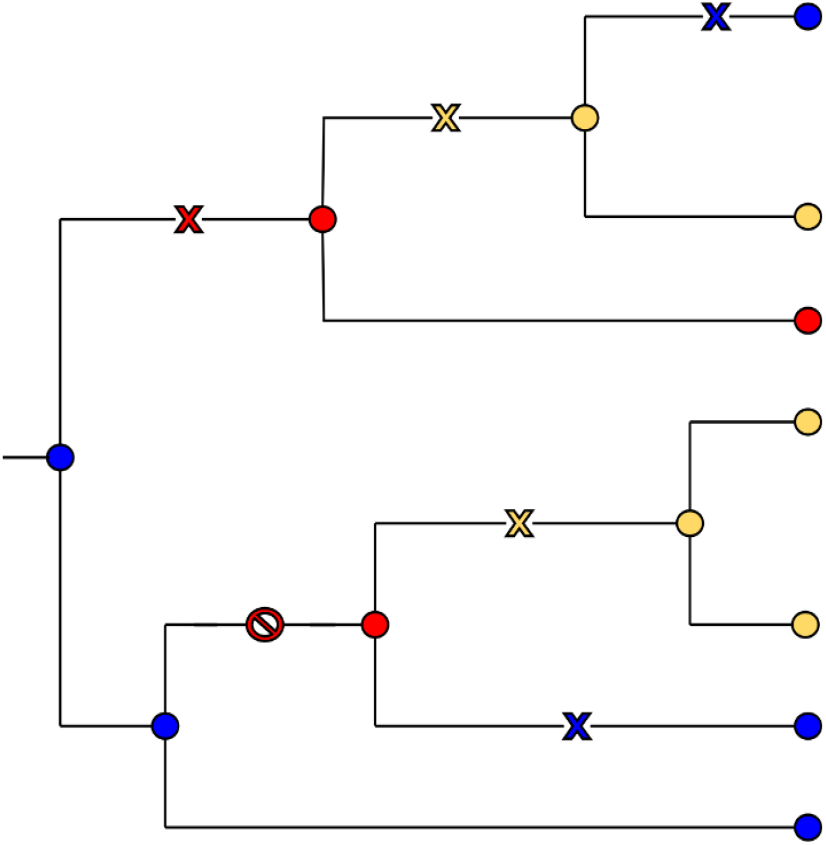
Counting the emergence events of mutations (EEMs). In this tree, we count two (parallel) EEMs toward the yellow state, two reversions toward the blue state, and only one EEM toward the red state since the mutation at the bottom of the tree is not transmitted to any leaf and thus not counted.

This way of counting EEMs has linear time complexity in the number of tips, just as the ACR and simulation algorithms, which explains the relatively fast computing times of the Emergence component (12 minutes on average per mutation on the rhodopsin dataset with 1,500 sequences, see below), though it is based on many (10,000) simulations.

### Correlation with the convergent phenotype

The correlation component of ConDor is based on the ‘Discrete’ method from BayesTraits (Pagel 1994; Pagel and Meade 2006), which combines Markovian modeling of trait evolution and Bayesian model comparison, to distinguish between the two hypotheses of independent (H_0_) versus dependent (H_1_) evolution of two traits along a phylogeny. Here, we apply BayesTraits to the analysis of two binary traits: presence/absence (1,0) of the mutation and convergent/not convergent (1,0) phenotype. For each of the hypotheses (corresponding to different evolutionary models), the marginal log-likelihood (approximated by the harmonic mean of the likelihoods after several millions of iterations) is calculated using a stepping stone sampler. BayesTraits then calculates the log Bayes factor (logBF) to decide if the (H_1_) dependence hypothesis is supported:

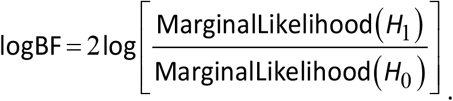

As described in the BayesTraits manual (www.evolution.reading.ac.uk/Files/BayesTraits-V1.0-Manual.pdf), a “logBF greater than 2 is considered as ‘positive’ evidence, greater than 5 is ‘strong’ and greater than 10 is ‘very strong’ evidence”. To account for different dataset sizes, we chose different thresholds for log BF (2 for the Sedges PEPC dataset, with 78 sequences, and 20 for the other datasets, with >1,000 sequences). Note, moreover, that when using the two components of ConDor, BayesTraits is only applied to mutations that are already selected by the Emergence component and corrected for multiple testing (regarding EEMs). Once we know that the evolution of the two traits is correlated, we need to determine the direction of the correlation: is the presence of the mutation favored (i.e. more frequent) in the convergent tips (positive correlation) or in the non-convergent tips (negative correlation)? For all the analyses presented here, we retained only positive correlations, corresponding to convergent molecular adaptations to the phenotype of interest (e.g. drug resistance in HIV). However, with the rhodopsin dataset, we were interested in adaptations to both environmental conditions (marine versus brackish/fresh water), and thus launched ConDor (and the other programs tested) twice, with each condition in turn considered “convergent”.

This method has been widely used in evolutionary biology and ecology to test correlations among behavioral, morphological, genetic and cultural characters, and for predicting functional gene linkages (Barker and Pagel 2005). To our knowledge, it has not been used to detect evolutionary convergence. One of the main advantages of this method is that it takes into account the phylogenetic correlation between taxa (as opposed to simple association tests, such as Fisher’s exact test that is commonly used for the detection of DRMs in HIV (Blassel et al. 2021)). Furthermore, it does not force the emergence of molecular convergence in all species with the convergent phenotype, as does the ‘One Change’ (OC) model of PCOC, for example (Rey et al. 2018). This characteristic is especially important as in most analyses we do not know the exact phenotype, but use a proxy. However, it should be kept in mind that the Correlation component in isolation can identify mutation events that fall outside the scope of convergent evolution. For example, a perfect correlation between a mutation and phenotype can arise from a single mutation event which is then propagated to all the tips of the corresponding subtree (a so called “founder” event; Bhattacharya et al. 2007; Gutierrez et al. 2019).

## Results

### Overview: data, methods and comparison criteria

We applied ConDor to three datasets with widely studied convergent mutations: (1) a sedge phosphoenolpyruvate carboxylase (PEPC) protein dataset with mutations associated with the acquisition of C4 metabolism; (2) an HIV dataset of reverse transcriptase with ∼33% sequences with drug resistance mutations (DRMs); and (3) a dataset of fish rhodopsin, a light-sensitive receptor protein that is highly conserved but known to vary at certain positions among species depending on their environment.

For HIV and rhodopsin data sets, we reconstructed the phylogeny from the sequences (nucleotide data and protein data respectively), using ModelFinder (Kalyaanamoorthy et al. 2017) and IQ-TREE (Nguyen et al. 2015) with standard options (see Material and Methods). For sedge PEPC data we used the provided phylogeny. Each phylogeny (with branch lengths reoptimized with amino-acid sequences for HIV and sedge PEPC) was used as input of ConDor, PCOC (Rey et al. 2018) and FADE (Murrell et al. 2012). For all tested methods, we evaluated the same mutations and positions, corresponding to the mutations present in at least 0.5% of the sequences and with more than 2 EEMs.

Given a rooted phylogeny, an alignment of amino acid sequences, and a list of convergent clades, PCOC performs a detection analysis for its three models (Profile Change, One Change, and both) for which we can set independent significance thresholds. Instead of detecting a change toward the same amino acid, the Profile Change (PC) component aims at detecting positions for which the general use in amino acid preference has changed in the convergent clades. This preference is modeled by a vector of amino-acid frequencies or ‘profile’ and, at a convergent position, the profile used in all convergent clades must be different from the ancestral profile. Conversely, for a non-convergent position, the same profile is used all along the tree. In addition, the One Change (OC) model forces that the switch of profile occurs along with at least one substitution in the branches rooting the convergent clades. Positions that verify the two sub-models are retained as convergent by PCOC, using a specific approach to combine the p-values from both sub-models. For the profiles, we used the C10 model that combines 10 profiles to represent the diversity of biochemical and mutational properties among amino acids (Le et al. 2008; default option in PCOC). Before running PCOC, users have to annotate the clades for their convergent status, using the list of species having the convergent phenotype. According to (Rey et al. 2018), a clade is said to be convergent if all its tips possess the convergent phenotype, and the branches yielding convergence (where OC is expected) are those rooting the maximal convergent clades. PCOC aims to detect positions with molecular convergence, and does not return a list of mutations, but a list of positions. We considered in our experiments that a mutation was detected by PCOC, when (1) it was present at a position where at least one of the sub-models (PC and/or OC) was verified (threshold above 0.8), and (2) the mutation in question was more frequent in species with the convergent phenotype than in the non-convergent ones.

FADE is one of the methods to detect selection available in the HyPhy package (Pond et al. 2005). FADE replaces previous approaches to test for episodic directional selection in protein alignments, which showed high detection power with DRMs in HIV (Murrell et al. 2012). To run FADE, the users first have to specify the branches that are expected to have undergone directional selection, called “foreground” branches. These typically correspond to all branches in convergent clades (and non-solely to the clade rooting branches, as with the OC component of PCOC; see Material and Methods for details). FADE tests for each position in the alignment if there is a “substitution bias toward a particular amino acid in the foreground branches, compared to the background branches”. The method relies on a Bayesian framework and a Bayes Factor >100 provides strong evidence that the site is evolving under directional selection.

ConDor aims at detecting mutations emerging more often than expected under a null model and which are correlated with the convergent phenotype (or its proxy). In our experiments, the null model corresponded to the best substitution model according to BIC, as inferred by ModelFinder (Kalyaanamoorthy et al. 2017). However, we also tested alternative models to check the robustness of the method to model violation (inevitable with real data). Both ConDor components use selection thresholds, which were fixed to corrected p-value <10% and log Bayes factor >20 for Emergence and Correlation respectively, unless otherwise specified. A mutation that verified both conditions was retained as a sign of foreground convergence.

We compared the three methods (PCOC, FADE and ConDor) using common statistics such as the number of true positives (TP: number of detected convergent mutations), true negatives (TN: number of non-detected non-convergent mutations), false positives (FP: number of detected non-convergent mutations) and false negatives (FN: number of non-detected convergent mutations). We also computed the recall (TP/(TP+FN)), precision (TP/(TP+FP)) and F1 score for each method. The F1 score is the harmonic mean of recall and precision:

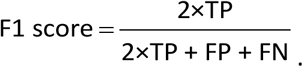

The F1 score provides a balanced view between recall and precision, which are generally in tension (improving precision typically reduces recall and vice versa). The F1 score is robust to class imbalance, as is usually the case with convergent mutations that are much less frequent that non-convergent mutations.

### Sedges PEPC protein dataset

We selected this dataset on C4 metabolism because it was the one used as a reference to evaluate PCOC in (Rey et al. 2018). This dataset comprises 78 sequences and allows comparison of convergence detection methods with a small dataset. C4 metabolism is a recognized case of convergence in plants and arose multiple times independently from the ancestral C3 metabolism. It is thought to be an adaptation to arid and warm environments (Ehleringer et al. 1997). Among the many proteins involved in the C4 photosynthetic pathway, phosphoenolpyruvate carboxylase (PEPC) has been studied to find a molecular basis for phenotypic convergence. PEPC is shared by both C3 and C4 plants and is encoded by a multigene family. The standard, well-supported hypothesis is that the *pepc* gene responsible for C4 metabolism has derived from an ancestral *pepc* gene responsible for C3 metabolism.

In our analyses, we focused on sedges, a plant family with multiple independent emergence events of C4 metabolism. We based our analysis on the dataset used in (Besnard et al. 2009) and later in (Rey et al. 2018). This dataset consists of an alignment of 78 sequences and 458 positions. We annotated the phenotype of the sequences according to two methods. First, based on the global phenotype of the plant (C3 or C4 metabolism) according to (Bruhl and Wilson 2007). With this annotation, multiple copies of PEPC in the same plant have the same annotation. Seven sequences were annotated to be intermediate between C3 and C4 and we removed this clade. This resulted in a dataset composed of 71 protein sequences, 22 being annotated as C4. Second, we kept the genotype-based annotation used by Besnard et al. (2009). This annotation is grounded in the fact that, in the PEPC amino-acid sequence, the A780S mutation (i.e., a change from A to S on reference position 780) has been experimentally demonstrated to be a major determinant of C4-specific characteristics. (Bläsing et al. 2000). They then predicted the metabolism associated with the sedges PEPC sequences according to the presence or absence of the A780S mutation. Compared to the first annotation, this resulted in a change of annotation for 4 sequences, from C4 to C3 metabolism. Moreover, 5 of the proteins from C3-C4 intermediate sedges were annotated as C4 and 2 as C3. This dataset, using the genotype-base annotation by Besnard et al. (2009), thus contains 78 sequences, 23 annotated as C4.

Mutations at positions 780 and 665 were confirmed experimentally to have an impact on the catalytic activity and folding of PEPC (Svensson et al. 2003; Christin et al. 2007). Including these two positions, Besnard et al. (2009) found 16 positions under positive selection that carry parallel amino-acid mutations in genes associated with C4 metabolism. Although most of these positions have not been fully confirmed experimentally, Rey et al. (2018) used these potentially convergent positions to evaluate the application of PCOC (and other approaches) to this dataset. We used the same reference positions in our analyses and comparisons and considered these positions to be true positives. We tested for convergence all mutations present in at least 3 sequences and with more than 2 EEMs for both phenotype- and genotype-based sequence annotations. Due to the difference in size of both datasets, we tested a different number of mutations. With the “phenotypic” annotation, we tested 59 mutations spread over 51 positions (with 11 positions likely harboring convergent mutations) and with the “genotypic” annotation, 66 mutations spread over 56 positions (with 12 positions likely harboring convergent mutations).

The results of the method comparisons for the two analyses are presented in Table 1. We first describe the results of the genotypic annotation as it was the one used in previous analyses (Besnard et al. 2009; Rey et al. 2018).

**Table 1:** Method comparison on sedge PEPC dataset. We display for PCOC, FADE, ConDor and their sub-components, several performance indicators on the detection of convergent positions either with genotypic annotation according to (Besnard et al. 2009) or phenotypic annotation. TP: true positives. FN: false negatives. FP: false positives. TN: true negatives. Recall (TP/(TP+FN)): proportion of TP among all positions retained by Besnard et al. (2009; 12 positions tested). Precision (TP/(TP+FP)): proportion of TP among all positions retained by the given method. F1 score: harmonic mean between recall and precision. PC: positions detected with a profile change in convergent clades, with posterior probability >0.8 (as used in (Rey et al. 2018)). OC: positions with one mutation on the branches leading to the convergent clades, with posterior probability >0.8. PCOC: combination of PC and OC components with posterior probability >0.8. FADE: positions with mutations showing evolution under directional selection in convergent clades, with Bayes Factor >100. Emergence: positions with mutations showing a number of EEMs statistically higher than expected with p-value <0.1 (after Holm-Bonferroni correction for multiple tests). Correlation: positions with mutations positively correlated with C4 annotation, with log Bayes factor >2. ConDor: combination of Emergence and Correlation. In bold: best result for each indicator.

| Genotypic annotation<br>based on A780S<br>56 positions tested | TP | FP | FN | TN | Recall | Precision | F1 score |
| --- | --- | --- | --- | --- | --- | --- | --- |
| PC | 1 | 4 | 11 | 40 | 0.08 | 0.20 | 0.12 |
| OC | 8 | 6 | 4 | 38 | 0.67 | 0.57 | 0.62 |
| PCOC | 7 | 3 | 5 | 41 | 0.58 | 0.70 | 0.64 |
| FADE | 11 | 4 | 1 | 40 | **0.92** | **0.73** | **0.81** |
| Emergence | 7 | 14 | 5 | 30 | 0.58 | 0.33 | 0.42 |
| Correlation | 7 | 6 | 5 | 38 | 0.58 | 0.54 | 0.56 |
| ConDor | 4 | 3 | 8 | 41 | 0.33 | 0.57 | 0.42 |
| Phenotypic annotation<br>51 positions tested |  |  |  |  |  |  |  |
| PC | 0 | 3 | 11 | 37 | 0 | 0 | 0 |
| OC | 0 | 1 | 11 | 39 | 0 | 0 | 0 |
| PCOC | 0 | 0 | 0 | 0 | 0 | 0 | 0 |
| FADE | 9 | 7 | 2 | 33 | **0.82** | 0.56 | **0.67** |
| Emergence | 6 | 17 | 5 | 23 | 0.55 | 0.26 | 0.35 |
| Correlation | 5 | 7 | 6 | 33 | 0.45 | 0.42 | 0.43 |
| ConDor | 4 | 3 | 7 | 37 | 0.36 | **0.57** | 0.44 |

Using PCOC on the genotypic dataset with a posterior probability threshold >0.8 (as used in (Rey et al. 2018)), 10 positions are detected among which 7 are true positives (TP). We thus find a large intersection between Besnard et al. and PCOC results, as previously described in (Rey et al. 2018). PCOC results are mostly driven by the OC component, which detects 14 positions including 8 TPs. The PC component, on the other hand, leads to lower accuracy and finds only 1 TP and 4 false positives (FP). With this dataset, PCOC results are derived primarily from the OC component that “assumes that convergent positions must have undergone a substitution on the branches where the adaptation took place” (Rey et al. 2019). With small datasets like this one, one can reasonably use PCOC recommended method, which infers “the branches where the adaptation took place” as the ones rooting the convergent clades, where all tips have the convergent phenotype. This works very well here (see Fig. 4 in Rey et al. 2018), hence the performance of PCOC. However, it is generally difficult (if not impossible) to define the position of these branches in larger and more complex phylogenies, due to phylogenetic uncertainty, reconstruction errors, and the use of a proxy for the phenotype. In this regard, with the phenotypic annotation PCOC and its subcomponents are no longer able to recover any TP. This shows that by annotating the sequences based on the genotype (presence or absence of the A780S mutation), there is a perfect match between the convergent clades and the mutations, which is advantageous for PCOC. However, on this data set, PCOC fails with the phenotypic annotation, which is mutation agnostic and probably more realistic for many convergent evolution studies.

Using the genotypic annotation, FADE detects 15 positions, including 11 TP. Even though this tool and model were designed in a different context (typically the detection of DRMs in viruses; Murrell et al. 2012), it performs very well and outperforms PCOC with a F1 score of 0.81 (against 0.64 for PCOC). With the phenotypic annotation, FADE accuracy decreases but still leads to the best results (F1 = 0.67) demonstrating the robustness of this method. This decrease is explained by the fact that there are fewer TPs recovered at the same time as more FPs. This behavior is expected as with this annotation, convergent mutations are no longer exclusively in convergent clades. Like PCOC, FADE requires the user to define the foreground branches where the adaptation occurred, which leads to similar difficulties. However, the hypotheses behind directional selection are less strict than with OC, as one simply assumes a mutational bias towards a certain amino acid in all branches of the convergent clades.

On this dataset, ConDor selected as null model the JTT substitution matrix associated with ‘freerate’ rates across-sites model (Susko et al. 2003; Soubrier et al. 2012) with 3 categories (R3). The Emergence component of ConDor detects 21 positions with higher-than-expected number of EEMs (Holm-Bonferroni adjusted p-value <10%), 7 of which are TP. Emergence does not use any phenotype information and likely detects convergent mutations linked to factors other than C4 metabolism, hence the high number of detected positions that do not belong to Besnard et al. list. Based on the genotypic annotation, the Correlation component refines these results, as expected since it accounts for the genotype and focuses on foreground convergent mutations: 7 of these 21 positions carry mutations that are positively correlated with C4 metabolism (BF>2; A780S, I588L, P540T, E572Q, S620C, H665N, F611L), among which 4 are TP. These correspond to the mutations present in the ‘ConDor detection zone’ in Figure 3. We notice that Correlation alone works fairly well (Table 1), without using any information on amino-acid exchangeability and biochemistry, as constitutive of the Emergence component. The combination of the two components in ConDor, using the genotypic annotation, increases the precision of the two components individually without however reaching the one of FADE and PCOC, resulting in mild F1 (0.42). However, F1 score overall decreases compared to the correlation component alone (0.56 for correlation alone). The 4 TPs found by ConDor are also found by the two other methods and especially the 2 positions that were demonstrated experimentally (780 and 665). All methods detect other convergent candidates, most of which being different between methods (see Fig. 3). This confirms that the experimental evidence on this dataset is still partial. Other convergent mutations could probably be found, and some of the 14 positions proposed by Besnard et al. might not actually be involved in C4 metabolism. With the phenotypic annotation, the F1 scores of the Emergence and Correlation components alone decrease as for the other methods (the Emergence component is not affected by the annotation, but less positions are tested than with the genotypic annotation). In contrast, ConDor as a whole is little affected by this annotation change (and the reduction of tests). It has the best precision among all methods (0.57) and maintains a mild F1 value (0.44). These results indicate that ConDor is robust to the detection of convergent mutations, even when a convergent mutation is not present in all the convergent clades or when a convergent clade also contains non-convergent mutations. Further analyses will confirm this finding.

**Figure 3:**
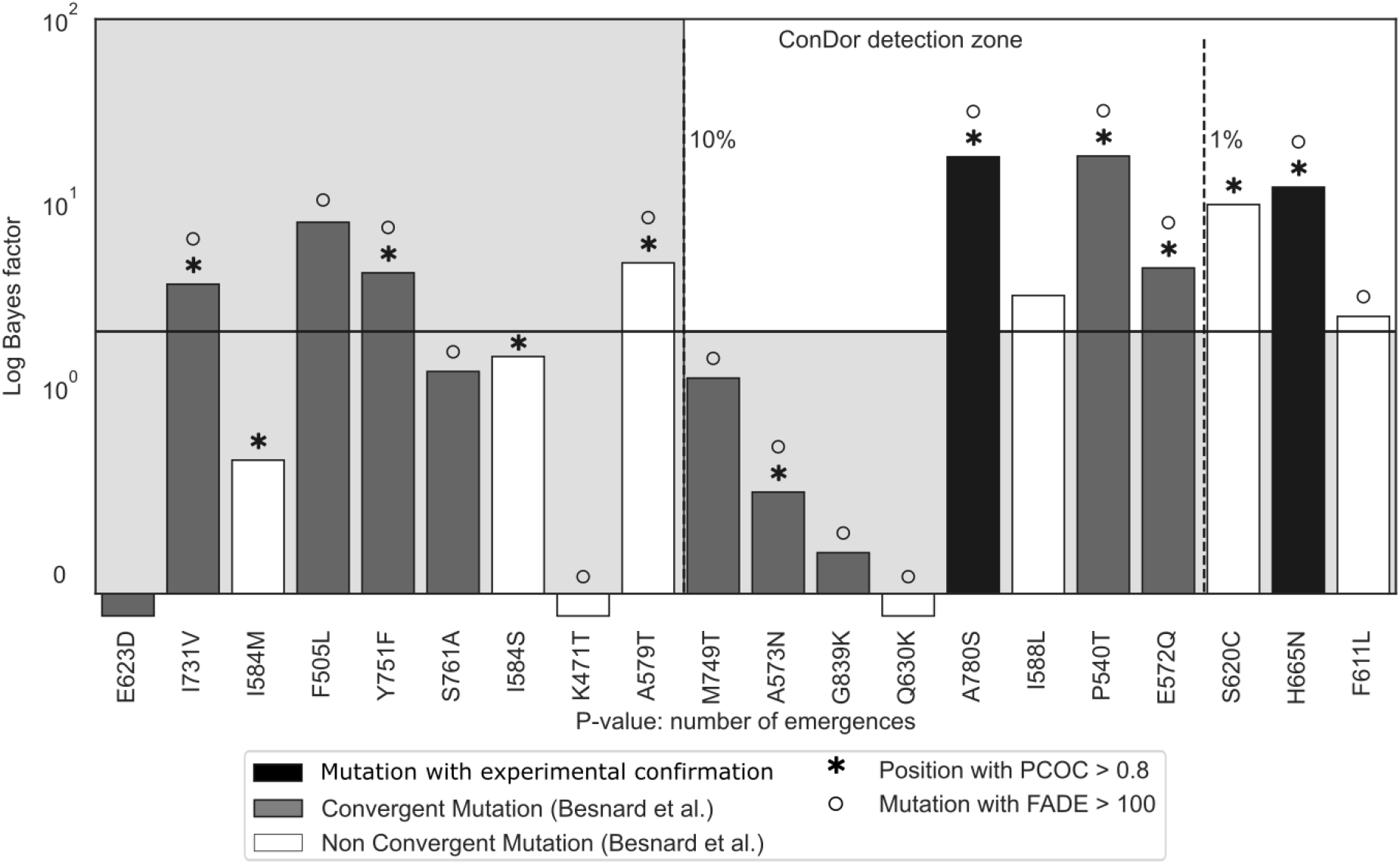
ConDor, PCOC and FADE convergent-mutation detections on sedges PEPC protein dataset. We display the mutations predicted to be associated to a C3-to-C4 change of metabolism by (Besnard et al. 2009). The two experimentally demonstrated mutations are in black, the other convergent mutations retained by Besnard et al. are in grey, and all other likely non-convergent mutations detected by PCOC, FADE or ConDor are in white. Mutations proposed by PCOC and FADE are indicated with an asterisk and a circle on the top, respectively. Mutations proposed by ConDor are present in the ‘ConDor detection zone’, corresponding to the upper-right white rectangle. Mutations are sorted on the x-axis by the p-value associated with the number of emergences (EEMs). The dashed lines represent various thresholds of Holm-Bonferroni adjusted p-values. We report on the y-axis the log Bayes Factor as obtained with BayesTraits. The plain horizontal line represents the threshold for ‘positive evidence’ of dependence between mutations and genotypic annotation (logBF >2).

### HIV reverse transcriptase dataset

Drug resistance mutations (DRMs) occur independently in patients undergoing drug therapy and are therefore a prime example of molecular convergence. In the case of HIV, they are well characterized and extensively studied, as their occurrence can lead to treatment failure and transmission of resistant viral strains. In particular, mutations must meet certain criteria to be identified as DRMs, including experimental validation (Wensing et al. 2019). DRMs are primarily found in proteins targeted by antiretroviral therapies: protease, reverse transcriptase, and integrase. The list of known DRMs affecting these proteins is publicly available at https://hivdb.stanford.edu/ and is updated regularly. As in the previous sedge PEPC dataset, DRMs are written in the form “XposY”, with X the ancestral (or wild-type) amino acid, “pos” the position of the substitution according to the HXB2 reference sequence, and Y the mutated amino acid, that is, the amino acid conferring resistance. We will use this notation for all our analyses.

In our case, we are interested in mutations on the reverse transcriptase, where DRMs are numerous, diverse, and have been experimentally confirmed. Furthermore, not all mutations occurring at a resistance-associated position make the virus resistant, but only a small subset, and frequently only one. This case is therefore particularly suitable for our method, which aims to detect convergence at the level of mutations and not only at the level of positions. Here we analyze an HIV-1 group M reverse transcriptase dataset sampled from 10 countries in West and Central Africa, and associated with metadata such as patient treatment status. We use treatment status as a proxy for phenotype, assuming that most patients with a detectable viral load (virus circulating in the host organism) are either treated patients whose treatment has failed due to the development of DRMs, or untreated (naive) patients without DRMs. However, some treated patients may have unsuppressed viral loads for other reasons (e.g., poor adherence to treatment), and some naive patients may have been infected with resistant strains harboring DRMs. This dataset was first studied in (Villabona-Arenas et al. 2016) and then in (Blassel et al. 2021), from which we retrieved the data. After removal of recombinant sequences (those for which the recombination occurs within the reverse transcriptase), it contains 1,858 sequences of 747 nucleotide positions that have been translated into 249 amino-acid positions. Ten subtypes and CRFs (circulating recombinant forms) are represented in this data, the major one being subtype C (37%). This dataset has several advantages for benchmarking convergence detection methods, compared to a UK dataset, also studied in (Blassel et al. 2021). First, a large percentage of the sequences are from treated patients (31%). Second, the DRMs are relatively frequent: ∼26% of the sequences harbor at least 1 DRM that is present in at least 10 sequences. Finally, there is relatively little transmitted resistance (12% of naïve sequences have one or more DRMs, Villabona-Arenas et al. 2016). For example, the mutation M184V, which is the most frequent, is observed in 378 and 5 sequences with treated and naive status, respectively, corresponding to relative frequencies of 66% and 0.4%. It is expected that such a DRM will be found by any reasonable convergence detection method. In contrast, rare DRMs are much more difficult to detect. For example, DRM Y188L is found in only 21 sequences (all from treated patients), which corresponds to 3.7% of treated sequences and 1% of all sequences. Note that in these two examples, we are far from observing a perfect correlation between the presence of the DRM and (the proxy used for) the phenotype (i.e., treatment status).

We tested 240 mutations present in at least 10 sequences and showing more than 2 EEMS, corresponding to 95 positions. Among these 240 mutations, 29 are DRMs distributed on 24 positions. We focused on these 29 DRMs to assess and compare the performance of our approach.

The PC component of PCOC works with a mild accuracy (F1 = 0.41). PC finds 12 positions associated with a shift in profile, 8 of which harbor DRMs. These 12 positions correspond to 20 mutations positively correlated with convergent phenotype, among which 10 are DRMs (TPs). In addition, no position is significant for the OC component (nor PCOC). This result is somewhat expected as DRMs are only found in a subset of all sequences showing the convergent phenotype. Moreover, several DRMs are not associated with a shift in profile and occur between closely related amino acids such as V and I or K and R (Fig. 4).

FADE with a default Bayes Factor threshold of 100 and with the HIVb substitution matrix has excellent recall but also detects many non-DRM mutations (66, Table 2), leading to a mild F1-score (0.38). Focusing on the detections with the highest BF (noted INFINITY in FADE’s outputs), FADE has a significantly higher F1 score (0.54). Overall, FADE’s performance on this dataset is good, which is consistent since the EDEPS and MEDS models (now replaced by FADE) were designed for drug resistance detection in HIV (Murrell et al. 2012). Running FADE with the JTT matrix (instead of HIVb), leads to similar recall, precision and F1 score, showing that FADE is robust to model misspecification.

**Table 2:**
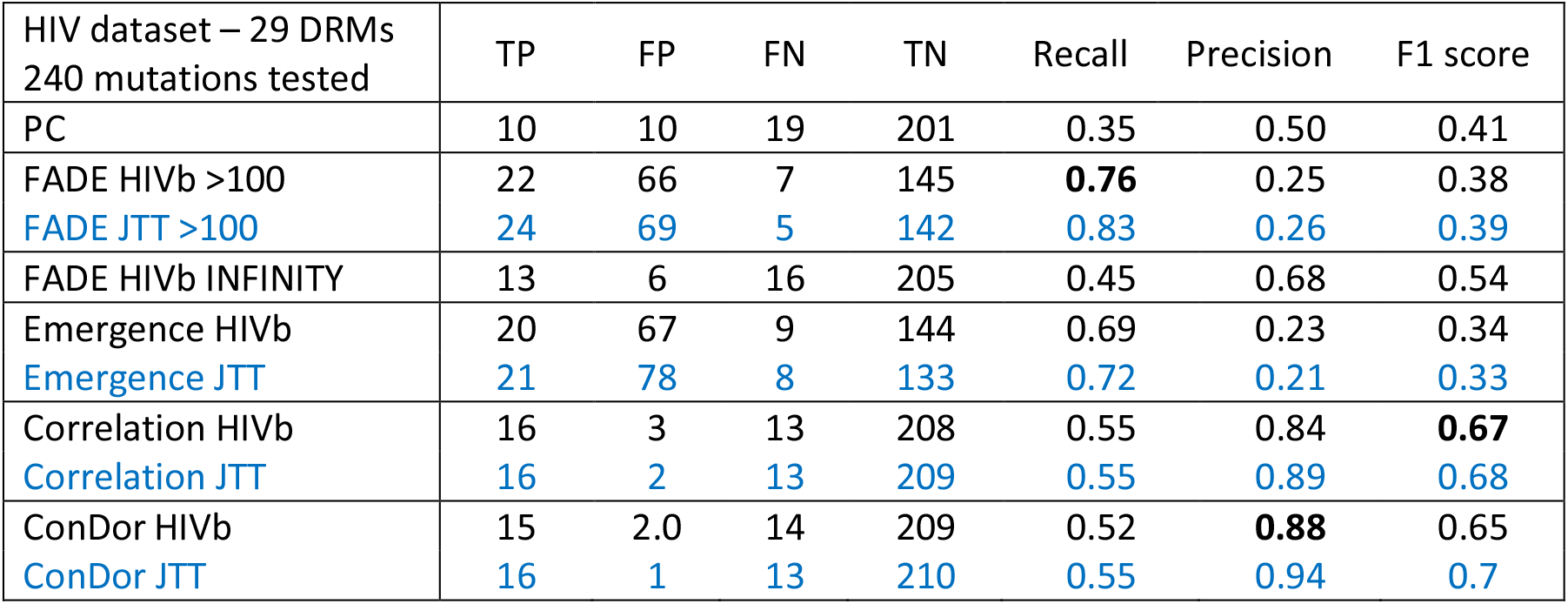
Method comparison on HIV reverse transcriptase dataset. Several performance indicators of the detection of convergent mutations are displayed for PC, FADE, ConDor, and ConDor sub-components. We display the results using the best substitution matrix (HIVb) and, when possible, with JTT that figures out a form of model misspecification. TP: DRMs found by the given method. FN: DRMs not found by the given method. FP: mutations found by the given method, which are not DRM. TN: non-DRM mutation, not found by the given method. Recall: proportion of TP among all DRMs (29 DRMs tested). Precision: proportion of TP among all mutations found by the given method. F1 score: harmonic mean of recall and precision. PC (profile change): mutations detected on positions with a profile change in convergent clades with a posterior probability >0.8 (Rey et al. 2018). FADE (INFINITY): mutations showing evolution under directional selection, with a Bayes Factor >100 (or INFINITY, i.e., >10^16^). Emergence: mutations showing a number of EEMs statistically higher than expected at a p-value <0.1 after Holm-Bonferroni correction for multiple testing. Correlation: mutations positively correlated with the treatment status, with a log Bayes factor >20. ConDor: combination of Emergence and Correlation. In bold: best result for each indicator, when using the best substitution matrix (HIVb).

The null substitution model inferred for this dataset using ModelFinder (Kalyaanamoorthy et al. 2017) is HIVb (Nickle et al. 2007), with ‘freerates’ (Soubrier et al. 2012) rates-across-site model and 4 rate categories. Using the Emergence component of ConDor, we detect 87 mutations with more EEMs than expected, after applying a Holm-Bonferroni correction for multiple testing (adjusted p-value <10%). Of these detections, 20 are DRMs, which represents a recall of 69% and is higher than expected by chance (Fisher’s exact test p-value = 2e-4). However, 67 mutations are non-DRMs events (column FP Table 2) and we do not know whether these are false positives or possible convergent candidates associated with phenotypes different from drug resistance (background mutations). The mutation-phenotype correlation, at a log Bayes factor greater than 20, detects 19 mutations including 16 DRMs (TPs). With this data, the Correlation component of ConDor is therefore sufficient, with similar results (slightly more sensitive, but slightly less precise) to those obtained using both components (F1 score = 0.67 vs 0.65 with ConDor; Table 2). As expected, the correlation between DRM and treatment status is strong, and treatment status is a good proxy for the resistance phenotype. We shall see in the following section that this configuration does not occur on the rhodopsin dataset, where both components are needed. With both ConDor components, 15 DRMs are detected as well as 2 mutations (T48S and L228R) that could be true convergent mutations. In particular, L228R (also detected by PC and FADE, Fig. 4) has previously been described as an accessory mutation occurring in response to certain HIV treatments (Rhee 2003; Blassel et al. 2021). In the case of model misspecification (illustrated by the JTT row in Table 2), the number of FP increases slightly, and lowers our precision (from 0.23 to 0.21). However, the Correlation component smooths this effect, showing that our method as a whole is robust to model misspecification. On this dataset, the Emergence component and FADE with BF >100 have similar performance in terms of accuracy and F1 score, even though the Emergence component does not have phenotype information. With both components, ConDor detects more DRMs and less non-DRMs (F1 score = 0.65) than FADE with INFINITY threshold (F1 score = 0.54). Both offer better accuracy than PC (F1 score = 0.41).

Regarding the ConDor false negatives (FN, undetected DRMs), we see (Fig. 4) that 5 of them are not detected because they do not pass the threshold limit of the log Bayes factor even though they have a significant p-value in terms of EEMs (K219E, M230L, V90I, E138A and P225H). However, 3 of them have a significant log Bayes factor (>10, K219E, V90I and P225H). Moreover, Sluis-Cremer et al. (2014) showed that FN E138A (with negative log BF) is a polymorphic mutation found naturally in naive patients and particularly in subtype C. This subtype happens to be the main subtype sampled in this dataset and mainly from naive patients (Villabona-Arenas et al. 2016). The false negative E138A is therefore prevalent in our dataset with no significant difference between treated and naïve patients, which explains our findings. Lastly, the false negative M230L is present in a small number of sequences (n=14) and the correlation with the phenotype is difficult to establish with such a small number. However, this DRM is significant for the Emergence component. There are 9 additional false negatives that were not detected by the Emergence component, 8 of which were also not detected by the Correlation component. Most of these mutations have a small number of EEMs and, as expected, both ConDor components here lack detection power.

**Figure 4:**
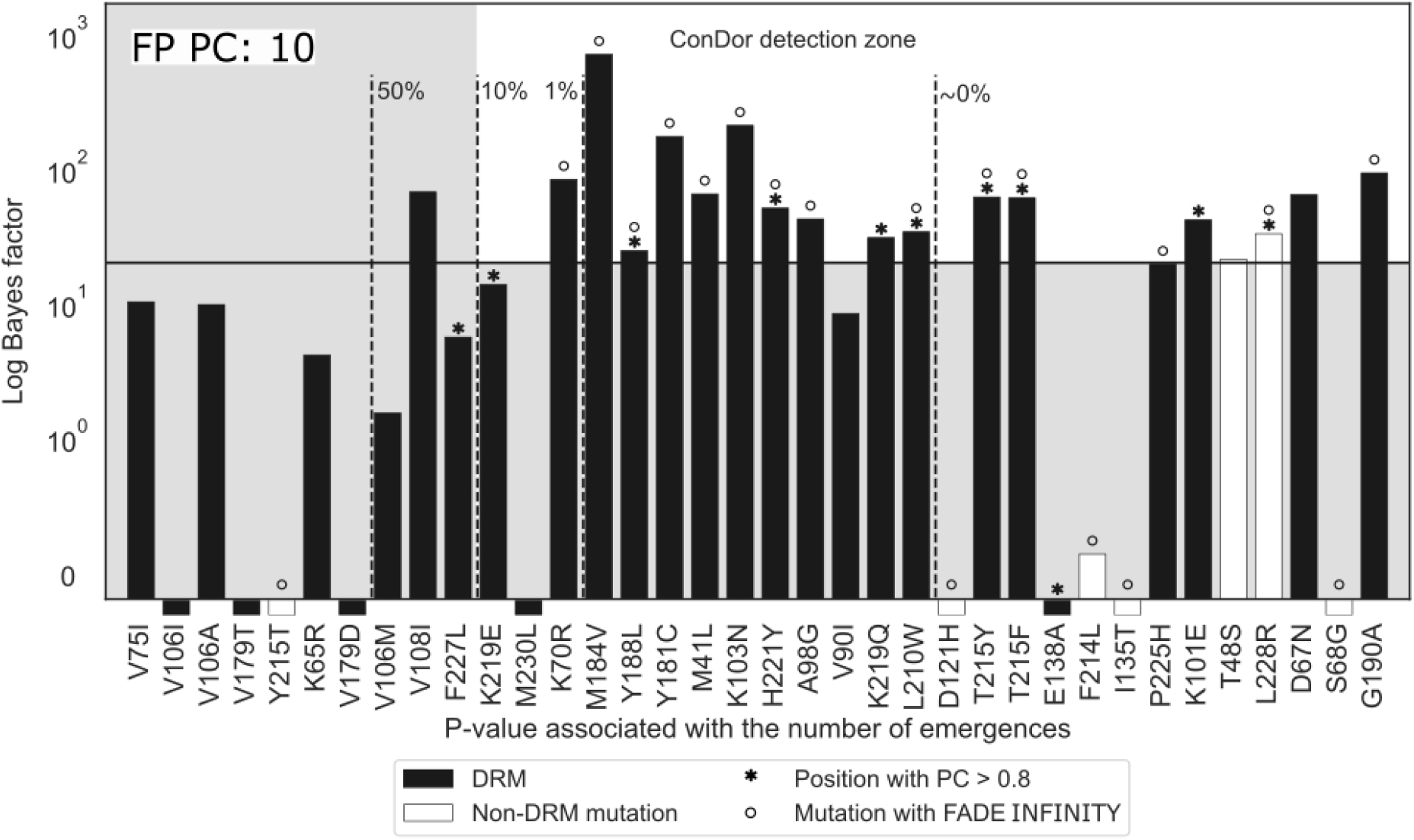
DRMs detection and convergent candidates on HIV data. We display all DRMs (black) and the non-DRM mutation (white) obtained using ConDor and FADE INFINITY on the HIV-1 group M reverse transcriptase dataset. If these mutations were found on positions associated with a shift in profile using PC, the bar is surmounted with an asterisk. If they were found associated with an INFINITY BF using FADE, the bar is surmounted with a circle. Mutations found by ConDor are present in the ‘ConDor detection zone’, corresponding to the upper-right white rectangle. Mutations are sorted by their p-value (Emergence component) on the x-axis. The dashed lines represent various thresholds of adjusted p-values using a Holm-Bonferroni correction. We report on the y-axis the log Bayes Factor as obtained with BayesTraits. The plain horizontal line represents the threshold for strong evidence of dependence between a mutation and the treatment status (log Bayes factor >20). Mutations that display a bar below the x-axis were found to be independent or negatively correlated with treatment status. FP PC in the upper-left indicates the number of false positives found with PC.

The Emergence component of ConDor is mostly driven by the number of EEMs and the exchangeability between amino acids. A mutation with a high number of observed EEMs, and corresponding to amino acids with low exchangeability, will rarely emerge in the simulations and will be detected by the Emergence component. Conversely, mutations between highly exchangeable amino acids, such as V and I, will often occur in simulations, which explains why DRM V108I is only detected by the Correlation component. This dataset contains several examples of DRMs involving highly exchangeable amino acids (some of which are TP detected by ConDor: K70R, M41L and, almost, V90I) demonstrating that convergent mutations with effects on phenotype can occur even between amino acids sharing highly similar biochemical properties. In this case, the PC component of PCOC may not be appropriate because it is designed to detect changes in amino acid profiles.

In this dataset, there is a strong correlation between most of the DRMs and the phenotype (treated/naive), hence the success of Correlation that has the best F1 score among all tested methods. In fact, with such HIV data, this genotype/phenotype correlation makes it possible to identify DRMs using simple association tests, with additional controls to account for the phylogenetic correlation between the sequences (Villabona-Arenas et al. 2016). We shall see that this is not the case for the Rhodopsin dataset where the proxy for the phenotype correlates less well with convergent mutations. In this case, both ConDor components are needed.

### Rhodopsin data

Rhodopsin is a photosensitive protein pigment responsible for the eye’s sensitivity to light. It is found in many vertebrates and has been shown to be under positive (or relaxed purifying) selection among species that evolve in different environments (Spady et al. 2005; Li and He 2009). Depending on the habitat and the amount of available light, different amino acids are observed at the same position, resulting in variations in structure of the rhodopsin and different maximum absorption wavelength (λmaxs). Mutagenesis experiments of engineered pigments revealed that the difference of λmaxs between most rhodopsins could be explained by 9 amino-acid mutations (Yokoyama 2008). In particular D83N, E122Q, F261Y and A292S (using similar substitution encoding as with HIV and PEPC) occurred several times independently and resulted into functional changes.

The dataset we used comes from a study in which the authors characterized substitution F261Y as convergent in fish rhodopsin, as a possible result of a transition from marine to brackish or fresh water environments (Hill et al. 2019). This dataset contains an alignment of 2,047 sequences with 308 amino-acid positions. The sequences have been classified by the authors into two groups: species found only in marine water and species that can live (exclusively or not) in brackish or fresh water. Some of the species associated with the habitat brackish/fresh water can therefore also be found in marine water. The proxy for the λmax is thus given by the environmental condition, depending on whether the fish species are found exclusively in marine water (43%) or not (57%). This approximation of the phenotype is rather imprecise, and we expect the correlation component to work less well on this dataset than for the sedge PEPC and HIV datasets. The reconstructed tree is well supported with 75% of the ultrafast bootstrap supports (Hoang et al. 2018) above 70%. We tested 358 substitutions, the ones present in at least 11 sequences and with more than 2 EEMs. In addition to D83N, E122Q, F261Y and A292S, the E122I and Y102F mutations have emerged several times in this dataset (Hill et al. 2019), and have been shown experimentally to affect absorption wavelength (Yokoyama 2008). We considered these mutations as well as their reversions as our true positives and explored their emergence for both habitats (marine and brackish/fresh water). Indeed, in the case of successive changes in the environment, we may find the reversion to the ancestral amino acid as convergent, when it emerged several times independently (i.e., N83D, Q122E, Y261F and S292A). All of this allowed us to study 10 mutations as truly convergent (F102Y and I122E did not appear 3 times in the data set). Because we were interested in adaptations to both the marine environment and brackish/fresh water, all programs were launched twice with each of the two conditions as the target.

We applied PC (profile change) and OC (one change) components of PCOC on this dataset for both environmental annotations. In total, 12 positions (corresponding to 27 mutations) were found to be associated with PC, 1 of which present mutations involved in a change of absorption wavelength of rhodopsin (E122I, E122Q and its reversion Q122E). This resulted in a low F1 score of 0.16 (Tab. 3). Moreover, as for the HIV dataset, no position was significant for OC and by extension for PCOC.

FADE with BF >100 showed a large number of detections, with 74 mutations under directional selection when the foreground branches lead to the taxa in brackish/fresh water and 55 mutations with marine water. This is hardly surprising given that the environment used as a proxy for the phenotype is very vague. A large proportion of the branches are labeled as foreground, which reduces the specificity of the method. Given this low specificity, mutations A292S, S292A, D83N, N83D, F261Y and E122I are detected by FADE, which has the best recall of all methods (0.6). However, FADE also has a low precision, resulting in a F1 score of 0.10, which is the lowest of all methods tested. With a more conservative threshold (BF >1000), FADE detects 40 and 49 mutations associated with marine and brackish/fresh environments, respectively, still including A292S, S292A, D83N, N83D, E122I and F261Y and resulting in a F1 score of 0.14. The F1 score decreases at higher thresholds. Similarly to the HIV dataset, FADE results are little affected by model misspecification (Tab. 3, FADE JTT).

The neutral model inferred by ModelFinder on this dataset was ‘MtZoa’ and ‘freerates’ with 8 rate categories. However, we analyzed the data using LG (second best substitution model) to ensure a fair comparison with FADE (MtZOA is not an available option). On this dataset, 60 mutations exhibit a number of EEMs significantly higher than expected as shown in Table 3. Using the Correlation component alone with a log Bayes factor of 20, one detects 73 mutations (40 correlated with brackish/fresh water and 33 with marine water) among which 6 are TP, resulting in a F1 score of 0.14. Combining both ConDor components, we find 18 convergent mutations that are correlated with the environment (9 with brackish/fresh water and 9 with marine water). Although predicting a few mutation candidates, ConDor still detects 4 out of the 10 convergent mutations experimentally confirmed by (Yokoyama 2008), which corresponds to the best F1 score (0.29) and precision (0.22) of all methods (Tab. 3). In case of model misspecification (Tab. 3, ConDor JTT), we see that the Emergence component is sensitive to this effect with more FP detected. However, ConDor is not affected by the change of model.

**Table 3:**
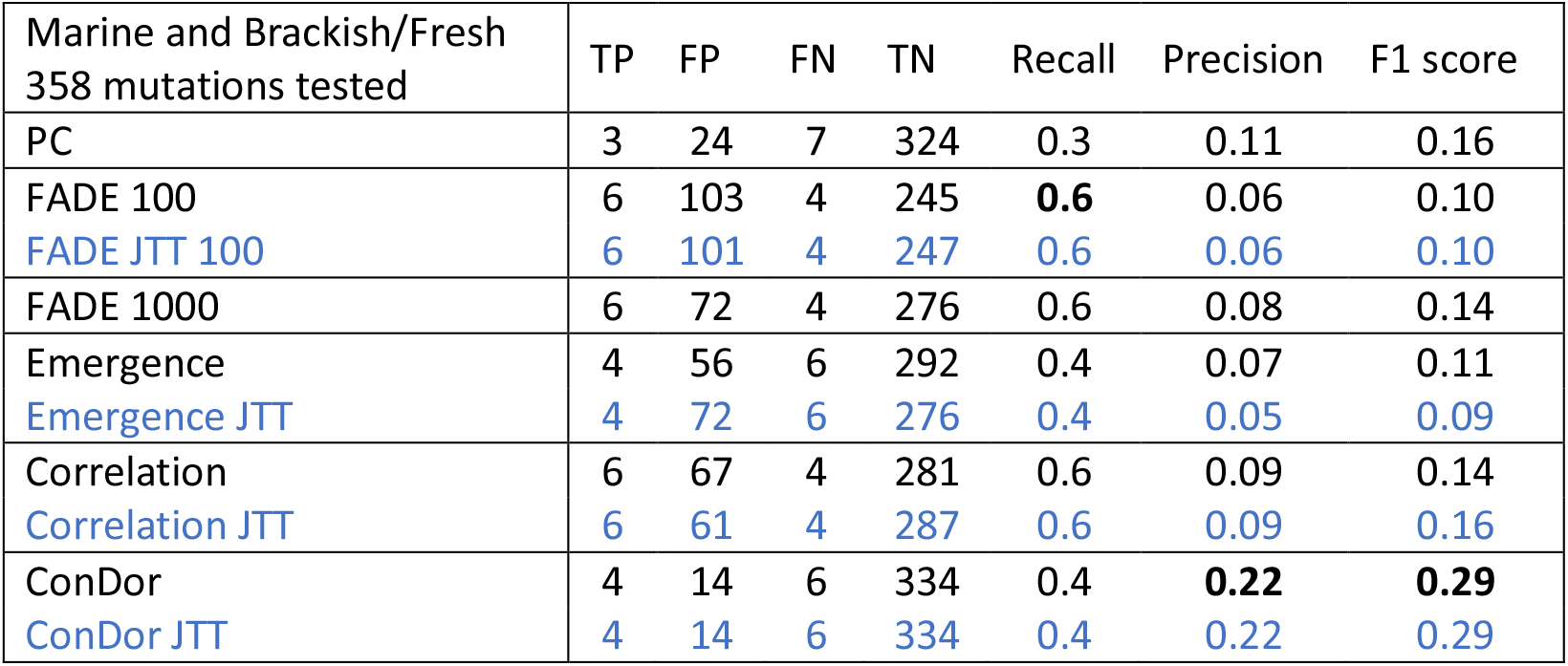
Method comparison on fish rhodopsin dataset. Several performance indicators on the detection of convergent mutations are displayed for PC, FADE, ConDor, and ConDor sub-components. We display the results using the second-best model (LG) for comparison purpose and, when possible, with JTT that figures out a form of model misspecification. TP: mutations affecting maximum absorption wavelength found by the given method. FN: mutations affecting maximum absorption wavelength not found by the given method. FP: mutations found by the given method, which are not experimentally demonstrated to affect absorption wavelength. TN: non detected mutations which do not affect absorption wavelength. Recall: proportion of TP among all mutations affecting maximum absorption wavelength (10 tested). Precision: proportion of TP among all mutations found by the given method. F1 score: harmonic mean between recall and precision. PC (profile change): mutations detected on positions with a profile change in convergent clades with a posterior probability >0.8 (Rey et al. 2018). FADE (1000): mutations showing evolution under directional selection, with a Bayes Factor >100 (or >1000). Emergence: mutations showing a number of EEMs statistically higher than expected at a p-value <0.1 after Holm-Bonferroni correction for multiple testing. Correlation: mutations positively correlated with the proxy of the phenotype, with a log Bayes factor >20. ConDor: combination of Emergence and Correlation. In bold: best result for each indicator.

As illustrated in Figure 5, ConDor retrieves mutations F261Y, D83N, A292S, and reversion N83D. E122Q is not found as convergent (adjusted p-value of ∼1 and log Bayes factor of 6 associated with the marine environment) because glutamine (Q) independently emerged only 3 times according to ACR, but emerged up to 11 times in simulations. Similarly, mutations E122I (3 EEMs) and Y102F (3EEMs) and reversions Q122E (3 EEMs), Y261F (3 EEMs) and S292A (11 EEMs) are not detected by ConDor as their number of EEMs was not significantly higher than expected. With this dataset, both components are needed to focus on a reasonable number of convergent candidates (PC and FADE with BF >1000 exhibit respectively 24 and 72 FP).

**Figure 5:**
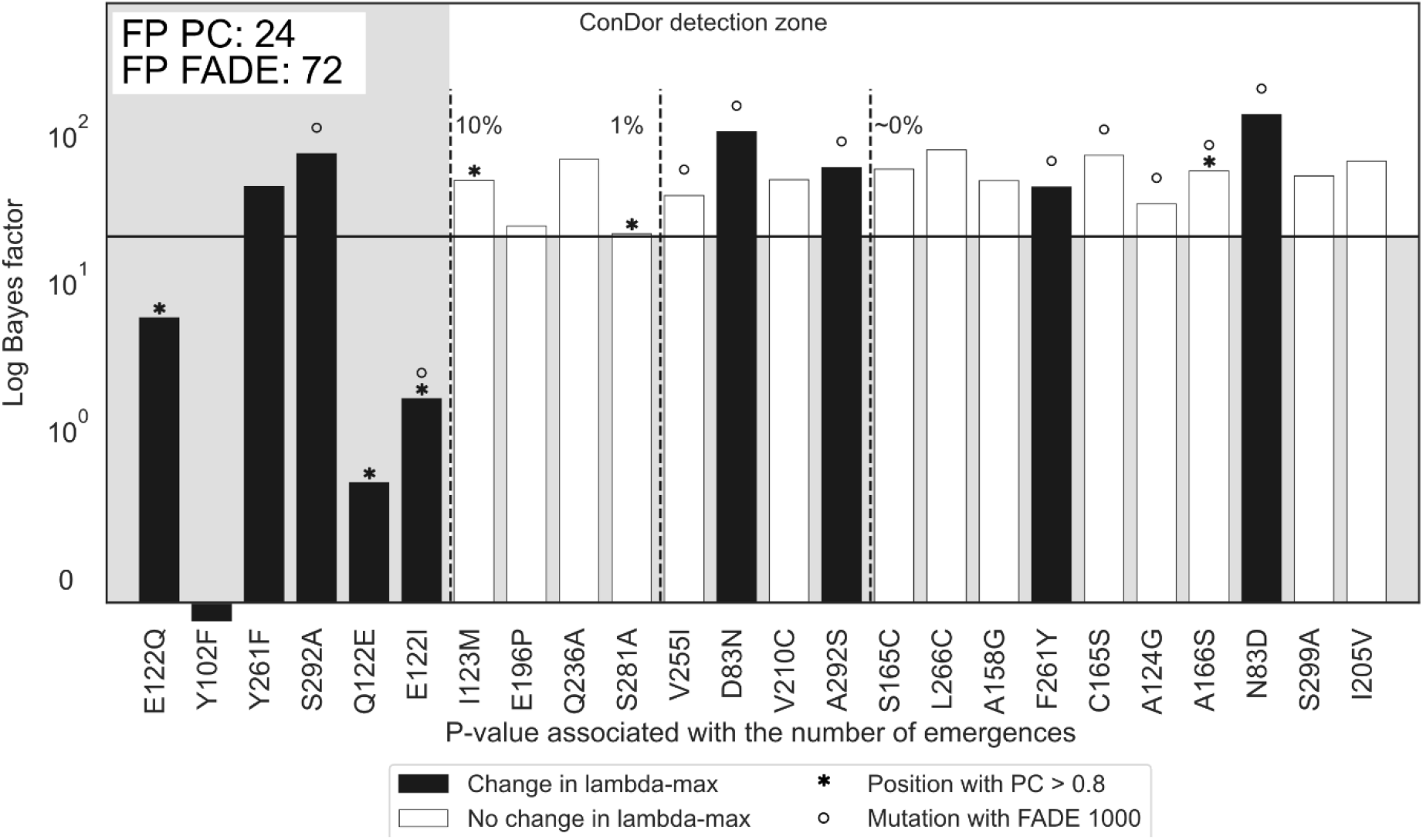
Detection of convergent mutations affecting maximum absorption wavelength on rhodopsin data. We display all mutations affecting maximum absorption wavelength (black) and the other detections (white) obtained using ConDor on the rhodopsin dataset. This figure combines both the detections correlated with brackish/fresh water and those correlated with marine water. If these mutations were found on positions associated with a shift in profile using PC, the bar is surmounted with an asterisk. If they were found associated with BF >1000 using FADE, the bar is surmounted with a circle. Mutations found by ConDor are present in the ‘ConDor detection zone’, corresponding to the upper-right white rectangle. Mutations are sorted by their p-value (Emergence component) on the x-axis. The dashed lines represent various thresholds of adjusted p-values using a Holm-Bonferroni correction. We report on the y-axis the log Bayes Factor as obtained with BayesTraits. The plain horizontal line represents the threshold for strong evidence of dependence between a mutation and the treatment status (log Bayes factor >20). Mutations that display a bar below the x-axis were found to be independent or negatively correlated with treatment status. FP PC in the upper-left indicates the number of false positives found with PC and FADE 1000.

Interestingly, convergent candidate mutation A166S is detected by the three methods (Fig. 5). This mutation is found by ACR to have 48 EEMs whereas on simulations it emerged at most 38 times. Following previous results from (Malinsky et al. 2015; O’Reilly et al. 2016), it might be associated with a blue-shifting absorption wavelength.

## Discussion

In this work, we developed the ConDor approach, which detects evolutionary convergence at the amino-acid mutation level using two components: Emergence and Correlation. Convergent (versus original) phenotypic annotations are given by users for extant taxa, without the need to define convergent clades and infer the phenotypic annotations of ancestral nodes. As we developed this method for the study of viruses and microorganisms for which the phenotype is difficult to access, ConDor allows the use of environmental conditions (or other selection pressures) as a proxy for the phenotype. Thus, convergent mutations can be found even if they are present in only a subset of the convergent taxa and if they are found in some taxa that do not possess the convergent phenotype. This is particularly suitable to the analysis of large datasets with several thousand sequences, where inference of convergent clades and ancestral phenotypes are especially challenging. For example, we were able to find more than half of the DRMs on a large HIV dataset where the application of PCOC was not appropriate because the underlying assumptions (OC and PC) were poorly satisfied. We also detected more DRMs with ConDor than using FADE while the assumptions of this software were made for DRM detection in HIV. Although it was primarily developed for the analysis of large datasets of viruses and microorganisms, ConDor was able to detect several convergent mutations involved in the change in metabolism in a small data set of sedge PEPC protein, and in the change in absorption wavelength in a large data set of fish rhodopsin. For the latter, its accuracy was markedly better than that of PCOC and FADE. These results confirm that ConDor detects a realistic signal of convergent molecular evolution and that it can be applied to a wide range of organisms and data sets.

We tested the robustness of the Emergence component of ConDor to model violation by using the JTT substitution matrix (Jones et al. 1992) instead of HIVb (Nickle et al. 2007) as the neutral evolutionary model for the HIV dataset study. A similar experiment was performed with fish rhodopsin where JTT was used again, instead of LG. In doing so, the sensitivity of Emergence remained high, we still detected the most frequent convergent mutations, but the number of false positives slightly increased. However, with the addition of the Correlation component, ConDor proved to be robust to these model violations. Since we never know the true evolutionary model, we expect that a substantial number of false positives may be observed with Emergence, even when using the best substitution matrix (as selected by IQ-TREE using BIC in our experiments). This behavior was observed in (Goldstein et al. 2015; Zou and Zhang 2015a), where the authors showed the difficulty to account for background convergent mutations using standard null models. More advanced substitution models based on CAT or CAT-JTT profile models (Lartillot and Philippe 2004; Le, Gascuel, et al. 2008), on mixture of matrix models accounting for structural features and rates of the positions (Le, Lartillot et al. 2008; Le et al. 2012), or on Markov modulated Markov models as in (Escalera-Zamudio et al. 2020), should likely improve the Emergence component, make simulations more realistic, and decrease the number of background convergent mutation detections in this component. However, these approaches are resource-intensive, and the Correlation component already complements the Emergence component well.

The two components of Condor can be used independently of each other. When using the Emergence component alone, there is no need to specify the taxa with the convergent phenotype. On the other hand, using the Correlation component alone, one loses the multiplicity constraint and cannot determine whether the detected correlation comes from a single founder event or from multiple independent emergences. Only the latter case corresponds to evolutionary convergence, which means that when using Correlation alone, we still have to perform an a posteriori check of the multiplicity of emergences.

ConDor was developed to detect convergent amino-acid mutations, not convergent positions, which makes it difficult to compare with existing approaches based on convergent position detection (e.g., PCOC (Rey et al. 2018)). An adaptation of ConDor to work at the position level could be an interesting feature to add to the program. Our approach is made possible because we work at the scale of a single protein with thousands of sequences, which provides sufficient signal and detection power. Working on thousands or even millions of positions (e.g., with bacterial genomes), ConDor would probably not have the statistical power to work at the scale of a single mutation due to multiple testing. An extension of ConDor to work at the gene level (similarly to (Chabrol et al. 2018)), or to detect convergence in a sliding window, would certainly be a useful development, allowing for the discovery of adaptive mutational patterns involving multiple sites in the protein alignment, rather than isolated sites as with the current version of ConDor.

Other improvements could concern the Correlation component that currently uses discrete trait evolution models in a Bayesian framework, which requires a lot of computing resources (∼30 min per mutation on the rhodopsin dataset). This computational burden could be greatly reduced using a similar maximum-likelihood approach (e.g., based on the ‘ace’ routine from APE; Paradis et al. 2004). In the same sense, simulations of the Emergence component are computationally expensive, and analytical approaches, inspired by to those used in (Chabrol et al. 2018), would also significantly reduce the computational burden of the approach.

## Materials and Methods

### Sedge PEPC protein dataset

Protein data of sedge PEPC, associated phylogeny, and “genotypic” C3/C4 annotation were downloaded from https://github.com/CarineRey/pcoc/tree/master/data/det. The protein data consists of a multiple sequence alignment of 79 protein sequences and 458 positions. The sequences are highly conserved, except for a few long deletions, and well aligned with no problematic gappy regions. We used the sequence *Chrysithr* as outgroup to root the tree and then removed it from the analysis (as Rey et al. 2018), resulting in an alignment of 78 protein sequences. Following (Besnard et al. 2009), 23 sequences have a convergent “genotypic” annotation (i.e. C4), based on the presence of the A780S mutation. For the “phenotypic” annotation, we annotated each gene using the annotation of the plant species it was sequenced from (Bruhl and Wilson 2007), and we removed the 7 genes from *Eleocharis baldwinii* and *Eleocharis vivipara* that perform both C3 and C4 metabolisms. This resulted in a multiple sequence alignment of 71 proteins and 458 positions, with 22 sequences annotated as convergent. The 7 sequences from *Eleocharis baldwinii* and *Eleocharis vivipara* were pruned from the provided phylogeny.

### HIV dataset

The HIV reverse transcriptase dataset we analyzed is based on the nucleotide alignment of (Villabona-Arenas et al. 2016), which was also studied in (Blassel et al. 2021), and is available from https://github.com/lucblassel/HIV-DRM-machine-learning/tree/main/data/African_dataset. This is a high quality alignment, thanks to the fact that the HIV proteins are highly conserved, with very few indels (see Villabona-Arenas et al. for details). This alignment consists of 3,990 HIV-1 group M partial reverse transcriptase sequences, divided into treatment-naive and treated sequences, along with a metadata file indicating the treatment status, whether the sequence has one or more DRMs and the subtype or CRF. The subtype annotation indicates that 2,247 sequences are recombinant forms, which we removed if the recombinant breakpoints were found within the reverse transcriptase (if so, we cannot reconstruct a sound phylogeny, n= 2,008). For example, 1,477 sequences were CRF02_AG, which recombines within the reverse transcriptase gene (Kusagawa et al. 2001). We then ran jpHMM (version of March 2015) (Schultz et al. 2012) to identify other possible recombinant forms. We used the default settings for HIV `-v HIV` and the priors provided in the jpHMM folder: `-a priors/emissionPriors_HIV.txt -b priors/transition_priors.txt.` Based on jpHMM analysis, we removed 124 additional recombinant sequences with breakpoints in the reverse transcriptase gene.

To root the tree, we added to this nucleotide MSA 3 reference sequences from the N group, which we downloaded from https://www.hiv.lanl.gov/content/sequence/NEWALIGN/align.html (reverse transcriptase: user-defined range 2550 – 3297).

The tree was inferred from the nucleotide MSA with IQ-TREE 1.6.8 while selecting the model (GTR+R10 in this case) with Model Finder (IQ-TREE option `-m MFP`). After rooting the tree, the 3 reference sequences of group N were removed from the analysis. The resulting alignment contains 1,858 group M sequences of 747 nucleotides that we translated into 249 amino acids. DRMs were identified in the translated MSA using the 2019 HIV-1 DRM list (Wensing et al. 2019). 571 sequences were annotated as treated and 1,287 sequences as naive. The tree branch lengths were re-optimized by ConDor using the protein MSA (see ConDor Pipeline Description below).

### Rhodopsin dataset

Rhodopsin protein data and fish habitat were downloaded from https://github.com/Clupeaharengus/rhodopsin/tree/master/phylogeny_habitat. We extracted from the “final_alignment.translated.fullrhodopsin.fasta“ alignment file 2,056 sequences corresponding to the identifiers indicated in the “spp_to_keep.txt“ file. After a quick tree reconstruction with FastTree (Price et al. 2009), we removed 7 badly aligned sequences (based on their aberrant branch lengths). We checked the quality of the resulting alignment with TCS (Chang et al. 2014) and obtained a score of 997/1000 demonstrating high reliability. Rhodopsin phylogeny was reconstructed from this protein MSA, using Model Finder (IQ-TREE 1.6.8 option `-m MFP`) to select the evolutionary model (MtZOA+R8) and IQ-TREE with option `--bb 1000` for ultrafast bootstrap approximation (Hoang et al. 2018). We rooted the tree using the same sequences as in (Hill et al. 2019) (*Huso huso* and *Polyodon spathula*) and removed them from the phylogeny for the analysis. This resulted in an alignment of 2,047 sequences, 883 annotated with marine water and 1,164 with brackish/fresh water. Habitat was provided in the “rabo_allele_hab.tsv” file from the repository provided in (Hill et al. 2019).

### PCOC

We used PCOC v1.0.1 (Rey et al. 2018) to detect convergent positions based on knowledge of genotype/phenotype (C3 vs C4), treatment status (treated vs naive), and habitat (marine vs brackish/fresh water). We used the C10 profile (`-CATX_est 10`) with 4 gamma categories `(--gamma`) and set the posterior probability threshold >0.8 for all models (PC, OC, PCOC) and data sets (`-f 0.8`). As described in the user guide (https://github.com/CarineRey/pcoc), the convergent scenario (`-m`) corresponds to the list of the maximal clades that exhibit the convergent phenotype. Each clade corresponds to an independent emergence event. Since it cannot be known exactly where the convergent transition occurred in the tree, the clades are first reconstructed by retrieving the tips with the convergent phenotype (C4 metabolism, treated, brackish/fresh water, marine water). Then, the internal nodes are recursively annotated with the convergent phenotype if the two child nodes also have the convergent phenotype.

### FADE

We used FADE 0.2 (unpublished to date) from the HyPhy package (Pond et al. 2005) to detect mutations under directional selection. FADE requires as input a rooted tree with annotations for the set of foreground branches suspected to have undergone directional selection. We annotated the foreground branches using http://phylotree.hyphy.org/. This software allows us to select terminal branches leading to tips with a convergent phenotype as foreground branches. Then, we can label internal nodes as foreground based on several methods (maximum parsimony, conjunction and disjunction). We labeled the internal nodes using conjunction, which is based on logical “AND” (a node is labeled foreground if all its children are foreground). This follows, in fact, the same labeling process as for PCOC. FADE was then run using the same substitution matrices as ConDor (JTT, HIVb, LG) and providing the same amino-acid alignments as for PCOC and ConDor.

### ConDor Pipeline Description

The ConDor Pipeline consists of several processes shown in Figure 1. Here, we describe the implementation details of IQ-TREE 1.6.8 (Nguyen et al. 2015) for the re-optimization step, PastML 1.9.33 (Ishikawa et al. 2019) for the ancestral reconstruction step, and BayesTaits 3.0.1 (Pagel 1994; Pagel & Meade 2006) for the correlation step.

#### Model selection, branch-length and evolutionary rate estimation

Given an input protein MSA and the corresponding phylogeny, we estimate the best evolutionary model and re-optimize the branch lengths and evolutionary rates for the protein MSA, using the `-m MFP` option of IQ-TREE, while fixing the topology using the `–te` option. This phylogeny with optimized branch lengths is used by ConDor, but also for PCOC and FADE analyses. In ConDor, site-specific evolutionary rates are retrieved from IQ-TREE with the `-wsr` option. For all analyses, we used the equilibrium frequencies corresponding to the substitution matrix, except for the model misspecification experiment with JTT on the HIV dataset, where we re-optimized the equilibrium frequencies (option `+FO`), in order to obtain a reasonable fit with the data, using a standard procedure. The best substitution model for each dataset was JTT+R3 (Sedge), HIVb+R4 (HIV) and MtZOA+R8 (Rhodospin). For the rhodopsin analysis with ConDor, we used LG+R8 to allow a fair comparison with FADE.

#### Ancestral character reconstruction by maximum likelihood

Ancestral character reconstruction in the ConDor pipeline is performed using PastML with option `--prediction_method MAP` (i.e. maximum a posteriori). PastML takes as input a parameter file (option `--parameters`) per position, containing (1) the amino acid frequencies for the entire alignment, and (2) a scaling factor for the position under study, corresponding to the evolutionary rate of the site, as estimated by IQ-TREE. This scaling factor (evolutionary rate) is used by PastML to re-scale the branch lengths while computing the tree likelihood. The selected substitution matrix (JTT, HIVb, LG) was given as input (`--rate_matrix`) using PastML option `-m CUSTOM_RATE`.

#### Correlation measurement using BayesTraits

Correlations between the convergent phenotype and mutations occurring more often than expected were measured using the BayesTraits ‘discrete dependent’ model. The convergent phenotype was annotated as 1 and the non-convergent phenotype as 0. Similarly, for a given position, the mutated amino acid of interest had value 1, and the other amino acids at that position had value 0. To assess whether dependence between the two traits was more likely than their independence, we followed http://www.evolution.rdg.ac.uk/BayesTraitsV3/Files/BayesTraitsV3.Manual.pdf. The dependence hypothesis was retained if the logarithm of the Bayes factor was greater than 2 for the sedge PEPC dataset and greater than 20 for the others. Thresholds of 2, 5, and 10 are given, respectively, as positive, strong, and very strong evidence against *H*_*0*_ in (Kass and Raftery 1995). Priors for transition rates were defined as uniform with a range between 0 and 100, as described in the user guide. Mutations that were found to be dependent of the phenotype by BayesTraits, were retained as convergent if the correlation was positive. To do this, we checked that the mutation frequency was greater in sequences with the convergent phenotype than in sequences with the non-convergent phenotype. More formally, let us denote: *m*_*C*_ the number of sequences that have the mutation M and are annotated with the convergent phenotype; *m*_*NC*_the number of sequences that also have the mutation M but are annotated with the non-convergent phenotype; *C* the total number of sequences annotated with the convergent phenotype; and *NC* the total number of sequences annotated with the non-convergent phenotype. If *m*_*C*_ /*C* > *m*_*NC*_/*NC*, then the correlation is positive, and M is considered as a convergent mutation by ConDor.

### Implementation

ConDor method is implemented in a Nextflow DSL1 v20.10.0 pipeline (Tommaso et al. 2017), taking as input an amino-acid alignment, a rooted tree, a file containing outgroup sequence identifiers and a file containing the list of sequences with the convergent phenotype. The python libraries `numpy` (Harris et al. 2020), `pandas` (McKinney 2010), and `scipy` (Virtanen et al. 2020) were used for data frames and matrices manipulations and for the statistics tools they provide. We used `biopython` (Cock et al. 2009) for sequence and alignment manipulations. Tree traversals and analyses were achieved with `ETE 3` (Huerta-Cepas et al. 2016). Graphs were obtained using the `matplotlib` (Hunter 2007) and `seaborn` libraries. All MSAs (translation to amino acids, subalignments, etc.) and tree manipulations (pruning, rooting, etc.) were performed using `goalign` and `gotree` (Lemoine and Gascuel 2021). Simulations and counting of EEMs were computed using homemade python scripts. Mutations emerging more frequently than expected were selected based on their p-value (with pseudo-count) after Holm-Bonferroni multiple testing correction, with an alpha risk of 0.1 This pipeline is available on Github at github.com/mariemorel/condor, via a webserver at condor.pasteur.cloud and as standalone in a docker. A user guide provides full details on the input and output formats, including explanations on the statistics provided for each mutation tested (p-value, Log-BF, genetic barrier, etc.).

## Data, results and tool availability

Our MSAs, phylogenetic trees, scripts and results analysis are accessible from the Github repository https://github.com/mariemorel/condor-analysis.

### Acknowledgments

We sincerely thank Luc Blassel, Jakub Voznica and Bastien Boussau for their help and suggestions.

